# Innovation and elaboration on the avian tree of life

**DOI:** 10.1101/2022.08.12.503188

**Authors:** Thomas Guillerme, Jen A. Bright, Christopher R. Cooney, Emma C. Hughes, Zoë K. Varley, Natalie Cooper, Andrew P. Beckerman, Gavin H. Thomas

## Abstract

Widely documented, megaevolutionary jumps in phenotypic diversity continue to perplex researchers because it remains unclear whether these dramatic changes can emerge from microevolutionary processes. Here we tackle this question using new approaches for modeling multivariate traits to evaluate the magnitude and distribution of elaboration and innovation in the evolution of bird beaks. We find that elaboration, evolution along the major axis of phenotypic change, is common at both macro- and megaevolutionary scales whereas innovation, evolution away from the major axis of phenotypic change, is more prominent at megaevolutionary scales. Indeed, the major axis of phenotypic change among species beak shapes at megaevolutionary scales is an emergent property of innovation across clades. Our analyses suggest that the reorientation of phenotypes via innovation is a ubiquitous route for divergence that can arise through gradual change alone, opening up new avenues for evolution to explore.

## 2 Introduction

Understanding patterns and constraints in the adaptive evolution of species traits is a major goal of evolutionary research. At macroevolutionary scales, renewed attention has been given to variation in the tempo and mode of trait evolution among lineages [1, 2, 3, 4, 5, 6, 7, 8]. This has led to growing recognition of the importance of major jumps or discontinuities in evolutionary rates as drivers of diversification of species traits [9, 10, 11]. For example, in both bird beaks and mammal skulls there are major jumps in the rate of trait evolution early in clade history as morphological diversity expands [9, 11] with lower magnitude jumps towards the present. Major jumps in species trait values are consistent with the concept of megaevolution, describing major discontinuities in phenotypes observed from the fossil record, first proposed by Simpson [12, 13].

Megaevolutionary dynamics appear to contrast with theory based on microevolutionary processes. Over microevolutionary timescales, phenotypic evolution is expected to follow lines of least resistance [14]. The term ‘line of least resistance’ has been used both specifically to describe phenotypic evolution along the axis of greatest genetic variation [14], and more loosely to describe the direction of phenotypic evolution with greatest observable variation [15]. Regardless of the definition used, the microevolutionary line of least resistance represents the direction with the most potential for evolution because it contains the most variation on which selection can act [16, 17, 18]. This expectation is supported by numerous studies showing that phenotypic divergence is often biased along (i.e. aligned with) the line of least resistance [15, 19, 20, 21, 22, 17, 23]. These studies suggest that alignment of phenotypic divergence with the line of least resistance may be common over comparatively short (1-2 million year) timescales. More generally, the alignment of macroevolutionary divergence with the microevolutionary line of least resistance is expected to decline over time [21]. However, there is evidence, for example from Anolis lizards, that stability in the direction of phenotypic evolution can extend over timescales spanning tens of millions of years [21].

The juxtaposition of microevolutionary predictions to macro- and mega-evolutionary observations is striking and raises the question of how phenotypic divergence accumulates across evolutionary scales. In particular, how can megaevolutionary jumps emerge from gradual micro- or macro-evolutionary processes of phenotypic evolution implied by microevolutionary predictions? [16, 18] The original descriptions of microevolutionary lines of least resistance explicitly refer to genetic constraints. However, at the macroevolutionary scale the major axis of phenotypic variation is an emergent property of broader set of genetic and developmental constraints interacting with selection on a moving adaptive landscape [24, 25]. At this scale the major axis of phenotypic variation among species [15, 26] can be estimated from a matrix of divergences in species mean phenotypic traits (e.g. the **D** matrix of [27, 21]. These matrices do not take into account phylogenetic relationships among species. In contrast, the **R** matrix of [28] describes the rate of among-species divergence in species traits and explicitly incorporates phylogeny (also see the **B** matrix of [29]) We hereafter refer to this **R** matrix as the *evolutionary rate matrix*. The major axis of the evolutionary rate matrix is a phylogenetic line of least resistance applicable at macro- and mega-evolutionary scales that is analogous to the genetic line of least resistance at microevolutionary scales.

By studying the phylogenetic line of least resistance and related properties of the evolutionary rate matrix [30, 31, 32] at different scales we aim to gain new insights into the macro- and mega-evolutionary trajectories that lead to the diversity of beak shapes among extant bird species and clades. While there is no strict definition that separates macro- and mega-evolution, we define the macroevolutionary scale as species divergence within clades and the megaevolutionary scale as divergence of clades within larger clades. The diversity of bird beaks has been well-studied at broad scales, particularly in relation to dietary ecology [33, 34, 35, 36, 37]. Previous studies have assessed how variation in rates of evolution of beak shape has generated this diversity [9, 38], revealing a general pattern of phylogenetically clustered evolutionary rates interspersed with major shifts deep in avian evolutionary history and in isolated lineages. Major shifts indicate lineages that occupy distinct areas of beak shape space relative to their close relatives. For example, a major shift in beak shape arises in the early divergence of Strisores separating hummingbirds (Trochilidae) from swifts (Apodidae), perhaps marking changes in ontogenetic trajectories [39] as hummingbirds diverged from broad billed ancestors (e.g. the Eocene stem Trochilidae fossils *Parargornis messelensis* and *Eocypselus rowei* [40, 41]). By assessing the multivariate directions of the evolution of beak shapes of species and clades we aim to deepen our understanding of how diversity in morphological form arises at macro- and mega-evolutionary scales in birds. To do this we use data on bird beaks derived from 3D scans of museum specimens for *>*8700 species (∼ 85% of all extant bird species) and generate evolutionary rate matrices describing the directions of multivariate beak shape evolution. The phylogenetic breadth of the data allows us to compare directions of beak evolution of subclades (e.g. orders) within the class Aves (the megaevolutionary scale) and of species within subclades (the macroevolutionary scale).

Various approaches have been developed to measure properties of trait matrices [42, 43, 44], broadly focused on describing the size, shape, and orientation of trait divergence [45]. Our specific focus is on divergence of species and clades along or away from the phylogenetic line of least resistance. We adopt concepts developed in the specific context of signal evolution to describe these alternative routes to divergence [46]. Specifically, Endler *et al*. (2005) refer to evolution of new phenotypes along the same major axis of phenotypic variation as *elaboration*. This is akin to phenotypic divergence among species following lines of least resistance, and, by definition, is expected to be a major driver of phenotypic evolution. In contrast, Endler *et al*. (2005) refer to evolution away from the major axis of phenotypic variation, either in random directions or along limited axes, as *innovation* [46]. Relative to elaboration, innovation is more likely to generate phenotypic novelty, but, if directions of phenotypic evolution are constrained, would not be expected to be a primary driver of phenotypic diversity among closely related species. We apply these terms to i) species divergence within clades, and ii) clade divergence within larger clades (Fig. 1). Both innovation and elaboration could in principle occur at the macro- or mega-evolutionary scale. We expect that elaboration is more important at macroevolutionary scales (species diverging within clades) and innovation more important at megaevolutionary scale (divergence among clades). For our bird beak data, macroevolutionary scales refer to the beak shapes of species elaborating or innovating relative to the phylogenetic line of least resistance of their taxonomic order, and megaevolutionary scales refer to avian orders elaborating or innovating relative to the phylogenetic line of least resistance for the entire class Aves. For clarity throughout the manuscript, we refer to elaboration and innovation as the general concept described above but use the following terms to distinguish macroevolutionary scale (species) and megaevolutionary scale (clade) elaboration and innovation: *elaboration*_*species*_, *innovation*_*species*_, *elaboration*_*clade*_ and *innovation*_*clade*_. We describe the algebraic formulation of elaboration and innovation for species and clades in detail in the Methods section. Using these novel statistical tools, we estimate the relative contribution of elaboration and innovation to the remarkable global radiation and diversity of bird beaks.

**Figure 1:**
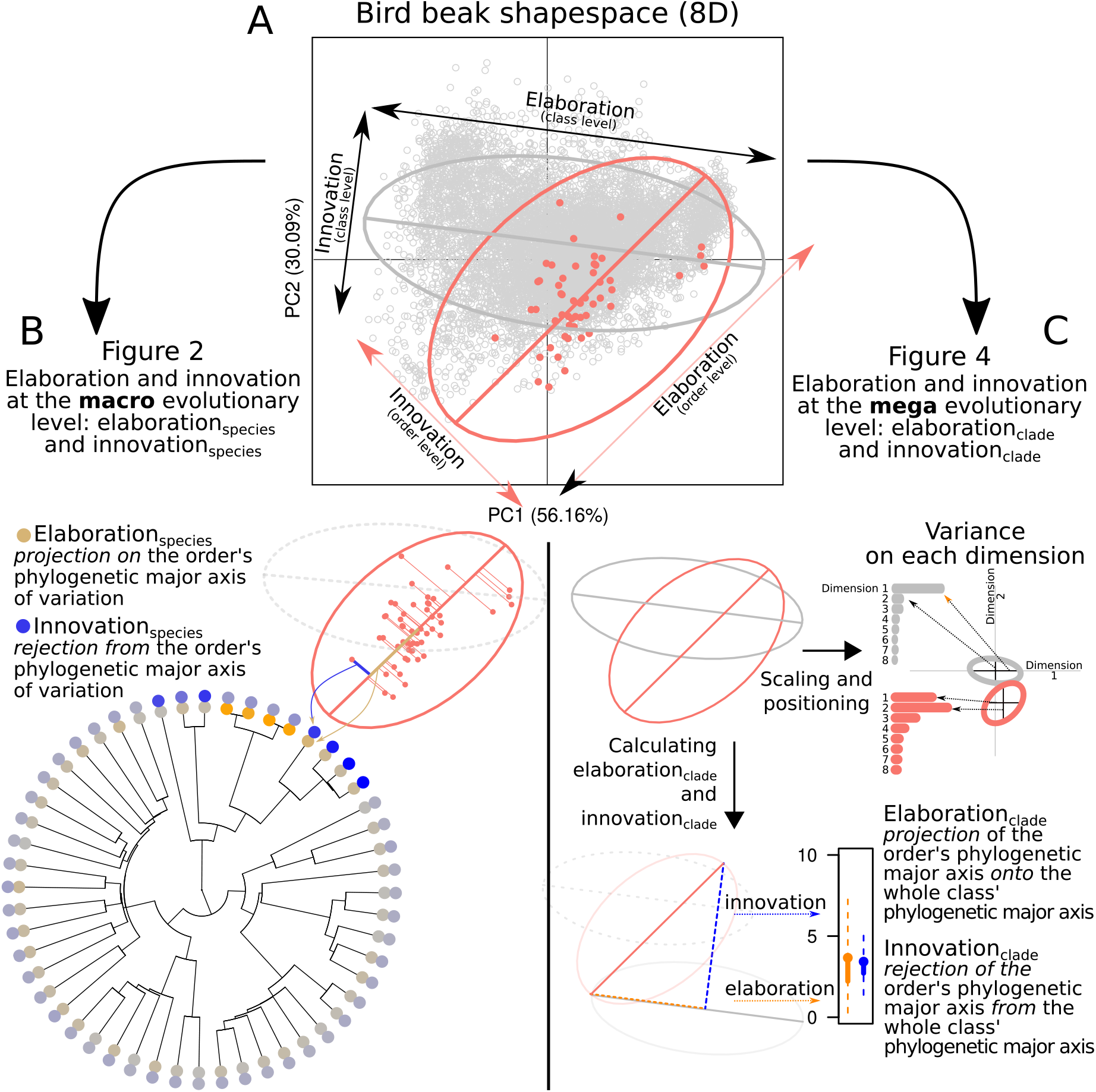
Relationships between key figures in the main text. A) represents the two first principal components (PCs) of the eight dimensional trait space with all bird beak shapes represented as gray circles and the Tinamiformes as red circles. The gray ellipse represents the overall phylogenetic major axis of beak variation in birds. Species aligned or away from this axis are respectively elaborators or innovators relative to the class Aves. The red ellipse and axis represents the phylogenetic major axis of beak variation in Tinamiformes only. Species aligned or away from this axis are respectively elaborators and innovators relative to the Tinamiformes order. B) In Figure 2, we measured elaboration and innovation at the macroevolutionary scale, i.e. the position of each species in terms of *elaboration*_*species*_ and *innovation*_*species*_ relative to their order’s phylogenetic major axis of beak variation. A species *elaboration*_*species*_ score is the species projection onto their order’s phylogenetic major axis of beak variation(i. e. their position on that axis) whereas their *innovation*_*species*_ score is their rejection from it (i.e. their distance away from that axis). The colours on the tips of the phylogeny correspond to the median *elaboration*_*species*_ (orange gradient) and *innovation*_*species*_ (blue gradient). C) Figure 4, we measured elaboration and innovation at the megaevolutionary scale, i.e. *elaboration*_*clade*_ and *innovation*_*clade*_. The ellipses are scaled and centered and represent the clade’s phylogenetic major axis of beak variation relative to the overall phylogenetic major axis of beak variation with the relative length of the ellipse on each dimension represented on the barplots as variance on each dimension. A clade’s *elaboration*_*clade*_ and *innovation*_*clade*_ scores are measured by projecting the clade’s phylogenetic major axis of beak variation onto the overall phylogenetic major axis of beak variation and measuring the projection (elaboration) and rejection (innovation) from this axis as described above. The distribution elaboration and innovation scores are taken from the projection of the 4000 pairs of evolutionary rate matrices with each pair being the focal level, e.g. order,and the parent level, e.g. the whole bird phylogeny.

## 3 Results and Discussion

### 3.1 Modeling nested trait covariance

Addressing the relative contributions of elaboration and innovation to the origins of biodiversity in deep-time requires: 1) large, multivariate datasets to allow exploration of trait covariances at different scales, 2) reliable and efficient computational methods to estimate the major axes of beak shape variation, and 3) a set of mathematical tools that can estimate degrees of elaboration and innovation at any scale. To meet the first challenge we use an eight-dimensional beak shape morphospace based on a geometric morphometric dataset of 8748 species of birds (described in [47]). To meet the other two challenges we introduce a novel analytical pipeline for measuring elaboration and innovation at the macroevolutionary scale (species within orders) and megaevolutionary scale (orders within the class Aves; Fig. 1 and S1). To estimate the major axis of beak shape variation we estimate the evolutionary rate matrix [28, 29] by fitting Bayesian phylogenetic generalised linear mixed model (pGLMM) models with beak shape as an eight-dimensional response variable [48]. The major axis can be identified from the leading eigenvector of the posterior distribution of evolutionary rate matrices. We regard the major axis of the evolutionary rate matrix as a deep-time analogue of the microevolutionary **G** matrix [49, 50] and posit that it represents the among species line of least evolutionary resistance, capturing the effects of historical contingency on multivariate evolution. This major axis of beak variation can be defined at any phylogenetic scale (i.e. for all species in the phylogeny, or for all species within any clade). We can then estimate elaboration and innovation based on that major axis: elaboration is defined as the position on the phylogenetic major axis of beak variation and innovation is defined as the distance from the phylogenetic major axis of beak variation (see Methods). To assess the roles of elaboration and innovation in beak shape evolution across scales we fit our pGLMMs as nested models in which we define evolutionary rate matrices, and therefore major axes of beak shape variation, for i) the entire phylogeny of all species included in the data (hereafter the class-wide phylogenetic level), ii) all species within each of nine clades mapping approximately to super-orders (the super-order level), and iii) all species within each of 27 clades mapping to taxonomic orders (the order level). These nested partitions of multivariate trait space provide the basis for all subsequent quantitative estimates of elaboration and innovation across scales.

### 3.2 Macroevolution: elaboration is a common route to divergence among species

We tested the expectation, derived from adaptive radiation theory [14, 21, 32], that species divergence is biased along phylogenetic lines of least resistance by calculating species-specific measures of *elaboration*_*species*_ and *innovation*_*species*_. If lines of least resistance are stable among species then we expect that species typically fall on, or close to, a conserved phylogenetic major axis of beak variation (*elaboration*_*species*_) rather than away from the axis (*innovation*_*species*_). To assess this we projected the beak shape data for each species onto the major axes of evolutionary rate matrices of their orders (see supplementary materials S5 for projection onto the major axes of beak shape variation from the class-wide phylogenetic major axis of beak variation or their super-order’s phylogenetic major axis of beak variation). We found typically higher values of *elaboration*_*species*_ than *innovation*_*species*_ (Fig. 2) and strong clustering of both *elaboration*_*species*_ and *innovation*_*species*_ (median *elaboration*_*species*_ for all orders = 0.114, 95%CI = 0.004-0.485; *elaboration*_*species*_ Pagel’s *λ* = 0.888; median *innovation*_*species*_ for all orders = 0.071, 95%CI = 0.018-0.224); *innovation*_*species*_ Pagel’s *λ* = 0.848).

**Figure 2:**
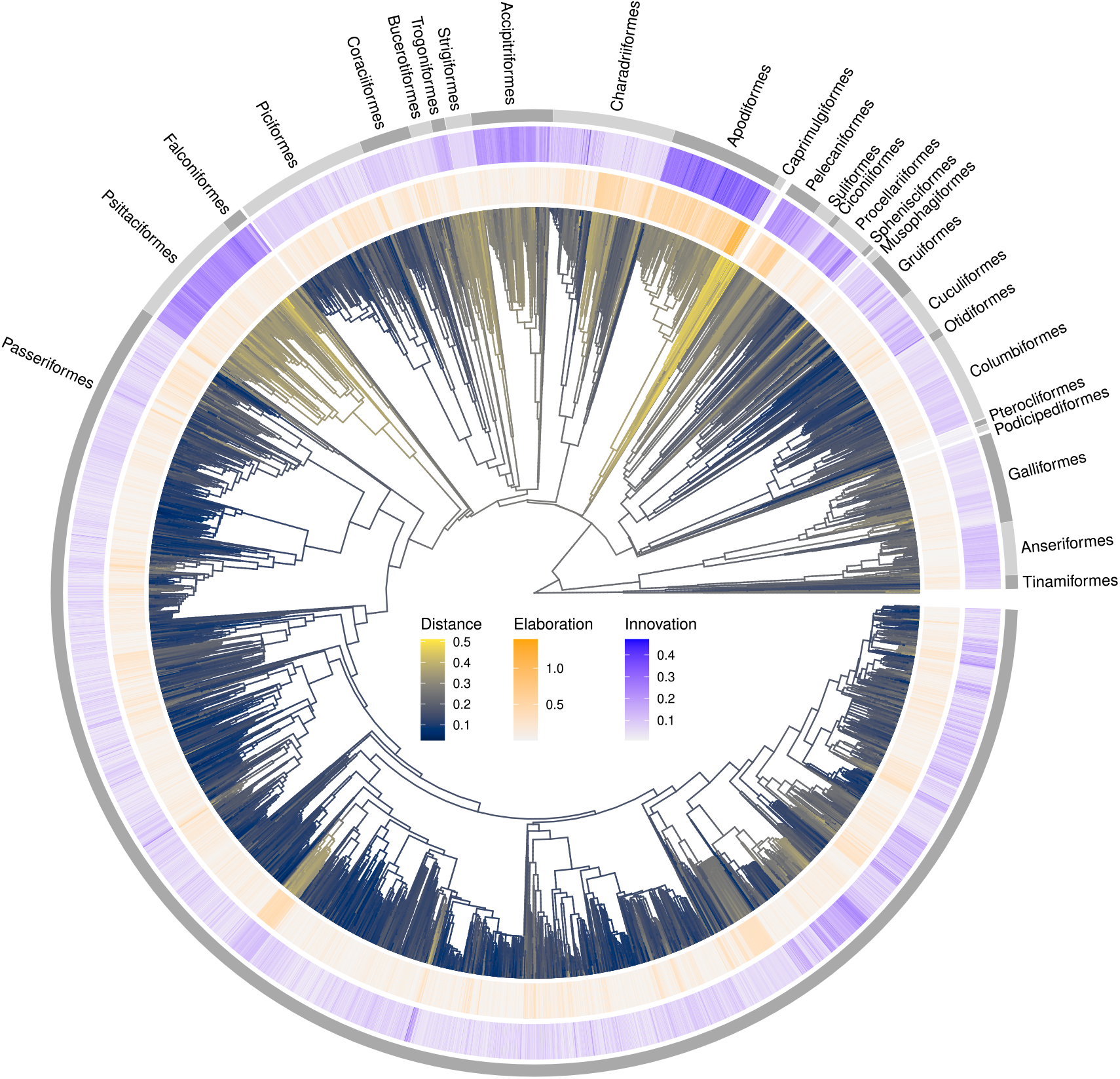
Avian phylogeny (n = 8748 species) showing Euclidean distance of species to the centroid of beak space (branches, cividis scale), and distributions of species beak shape *elaboration*_*species*_ (inner circle, orange scale), and *innovation*_*species*_ (outer circle, blue scale). Elaboration and innovation scores represent comparisons of species at the order level. Additional comparisons to super-orders and at the class-wide phylogenetic level are shown in Fig. S5 and for family and super-family comparisons of the order Passeriformes in Fig. S14.

*Innovation*_*species*_ is more apparent within non-passerine, rather than passerine, clades. Most notably we see higher levels of innovation in orders with hook-shaped beaks (Pstittaciformes, Falconiformes, Accipitriformes), and the Apodiformes (swifts and hummingbirds). Clades with hooked beaks have highly integrated beaks and brain cases and strong allometric patterns of shape variation, particularly among raptorial clades [35, 51]. Apodiformes are also notable for having both high innovation and elaboration scores. The presence of extremes of elaboration and innovation might reflect the deep divergence and distinct ecologies of the two families, Trochilidae (hummingbirds) and Apodidae (swifts), that constitute the order. The major axes of beak divergence within the order follow a trend from swifts (short, wide-gaped beaks; aerial insectivores) to hummingbirds (long, narrow, and often curved beaks; primarily nectarivores) that is not present within either family. This phylogenetic major axis of beak variation is likely to be representative of innovation between clades, rather than a line of least resistance within a clade.

Phylogenetic distributions of *elaboration*_*species*_ and *innovation*_*species*_ broadly hold whether comparisons are made at the order, super-order (*elaboration*_*species*_ Pagel’s *λ* = 0.853; *innovation*_*species*_ Pagel’s *λ* = 0.914), or class-wide phylogenetic level (*elaboration*_*species*_ Pagel’s *λ* = 0.935; *innovation*_*species*_ Pagel’s *λ* = 0.837; Fig. S5 and S6) and in further nested analysis of the order Passeriformes (Fig. S15 and S14). The extent of both elaboration and innovation tends to reduce as we move from class-wide phylogenetic to within-order comparisons. We regard within-order comparisons as the most informative level of comparison at the macroevolutionary scale because comparisons of species level values derived from the main axis of variation at the level of whole Neornithes might reflect global constraints in beak shape or an axis of innovation among clades (as with the Apodiformes discussed above). The class-wide phylogenetic major axis of beak variation aligns closely with the raw (non-phylogenetic) major axis in the trait space and primarily describes variation between short, deep, and wide beaks at one extreme, and long, shallow, and narrow beaks at the other. Our results point to the possibility that within clades the phylogenetic major axes of beak variation follow different directions.

Collectively, our results suggest that elaboration is the most common route by which species beak shape evolves. While *elaboration*_*species*_ is the main mode of beak shape divergence at the macroevolutionary scale (Fig. 2), it is also commonly positively correlated with *innovation*_*species*_ (Fig. 3 and S8). In 14/27 clades we find significant positive correlations with non-significant but positive trends in a further 11 clades. Only two clades (Falconiformes and Columbiformes) hint at a negative correlation but are clearly non-significant. The apparent predominance of *elaboration*_*species*_ when measured at the macroevolutionary scale is consistent with the idea of evolution along lines of least resistance yet the lack of trade-off between *elaboration*_*species*_ and *innovation*_*species*_ further suggests that evolvability, which is measured by elaboration along a single axis, may in fact be better regarded as multi-dimensional. Indeed, if divergence can arise simultaneously along multiple axes then this may help to resolve the paradox of apparent shifts in phenotype that have contributed to the diversity of species morphologies across the tree of life.

**Figure 3:**
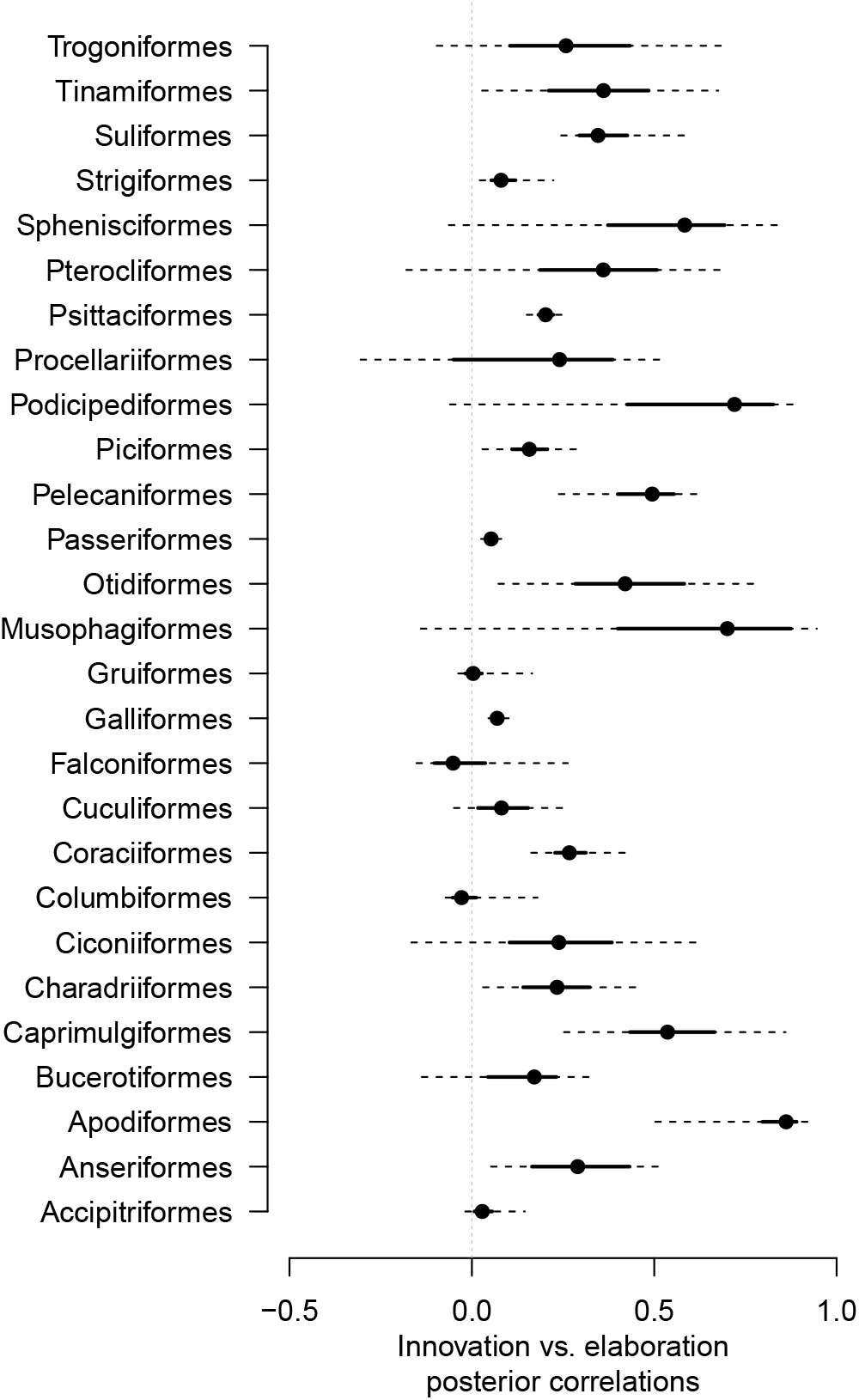
Posterior correlations between *elaboration*_*species*_ and *innovation*_*species*_ for each super-order. The dots, thick lines and dashed lines represent respectively the median, 50% confidence interval (CI) and 95% CI scores. 14 out of 27 of the orders have a clear posterior correlation between *elaboration*_*species*_ and *innovation*_*species*_ (i.e. 95% CI does not overlap with 0), nine have a somewhat positive posterior correlation (i.e. 50% CI does not overlap with 0) and four have no clear posterior correlation. Note that no order has a clear negative posterior correlation. This trend suggests that there is no tradeoff between *elaboration*_*species*_ and *innovation*_*species*_ and that both can be common routes for beak shape evolution at the macroevolutionary scale.

### 3.3 Megaevolution: multiple routes to innovation throughout avian evolutionary history

If deep-time jumps in beak shape [9] are the result of innovation we would expect to see changes in the orientation of trait space of clades (i.e. orders or super-orders) relative to the phylogenetic major axis of beak variation of the whole class Aves. To test this, we measured the *elaboration*_*clade*_ and *innovation*_*clade*_ by translating the phylogenetic major axis of beak variation of each clade’s (i.e. super-orders and orders) evolutionary rate matrix onto the class-wide phylogenetic major axis of beak variation so that they shared the same origin in the shapespace. We then used linear algebra to project a focal clade’s phylogenetic major axis of beak variation onto the class-wide phylogenetic major axis of beak variation (Fig. 1). We interpret the measured linear algebraic projection (Euclidean distance along the phylogenetic major axis of beak variation) as the clade’s *elaboration*_*clade*_ score and the measured linear algebraic rejection (Euclidean distance from the phylogenetic major axis of beak variation) as the clade’s *innovation*_*clade*_ score. A high *innovation*_*clade*_ score indicates that the direction of evolution of a clade differs from the direction of its parent clade (e.g. the direction of evolution of an order differs from its parent super-order). These scores were calculated for each of 4000 posterior pairs of clade *vs*. evolutionary rate matrix (Fig. 4 blue and orange lines). We found that for most clades the phylogenetic major axis of beak variation is not aligned with the class-wide phylogenetic major axis of beak variation (Fig. 4). Despite the limitations of representing eight-dimensional space in 2D, our plots of the average elliptical representation of the evolutionary rate matrix for the class-wide phylogenetic major axis of beak variation and each super-order and order (Fig. 4) highlight the striking variation in the orientation of beak shape major axes among clades (Fig. 5). More than half of the assessed clades (4/8 super-orders and 15/27 orders) displayed higher median *innovation*_*clade*_ than median *elaboration*_*clade*_ scores implying that *innovation*_*clade*_ is a more common generator of beak shape diversity than *elaboration*_*clade*_ in deep-time (i.e evidence at the clade scale).

**Figure 4:**
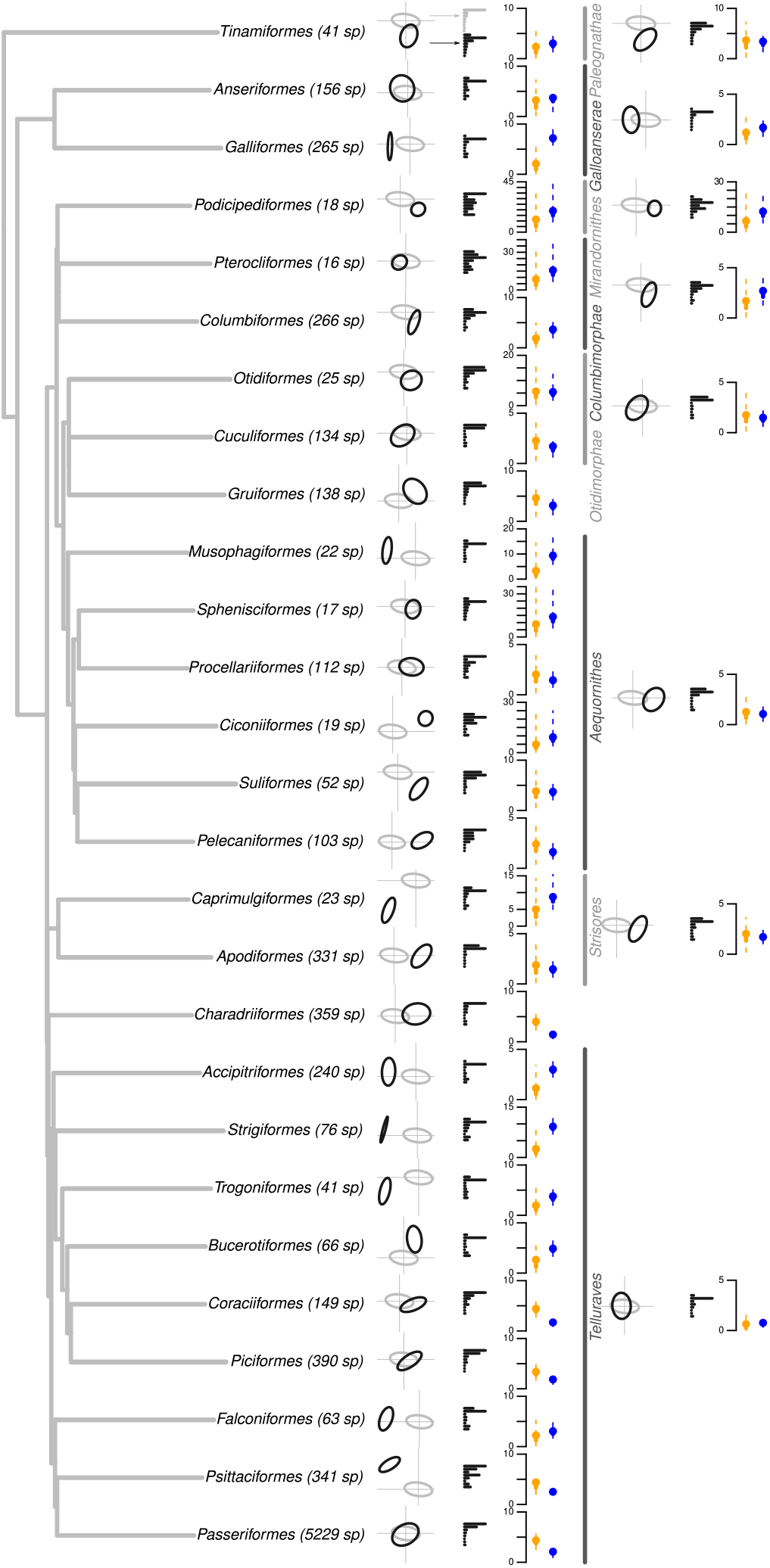
*Elaboration*_*clade*_ and *innovation*_*clade*_ at the megaevolutionary scale for each order and super-order in the bird phylogeny. The black and gray ellipses are the scaled average evolutionary rate matrices from the pGLMM models for respectively the clades (black) and the class-wide phylogeny (gray). The ellipses are centered on the position of the clade in the shapespace. The associated black bar plots represent the variance on each of the eight dimensions in the shapespace with the top gray bar plot representing the variance on each of the eight dimensions for the class-wide phylogeny. The orange and blue distributions represent respectively the distribution of the *elaboration*_*clade*_ (orange) and the *innovation*_*clade*_ (blue) scores for each clade and the dots are the median *elaboration*_*clade*_ or *innovation*_*clade*_ and the solid and the dashed lines representing the 50% and the 95% confidence intervales of the distribution of the scores across the 4000 posterior samples. The ticks on the y-axis always represent five arbitrary units of elaboration and innovation. An alternative visualization of the same information is available in the supplementary materials in Fig. S12 along with a companion plot showing further nested structure within the Passeriformes in Fig. S15.

**Figure 5:**
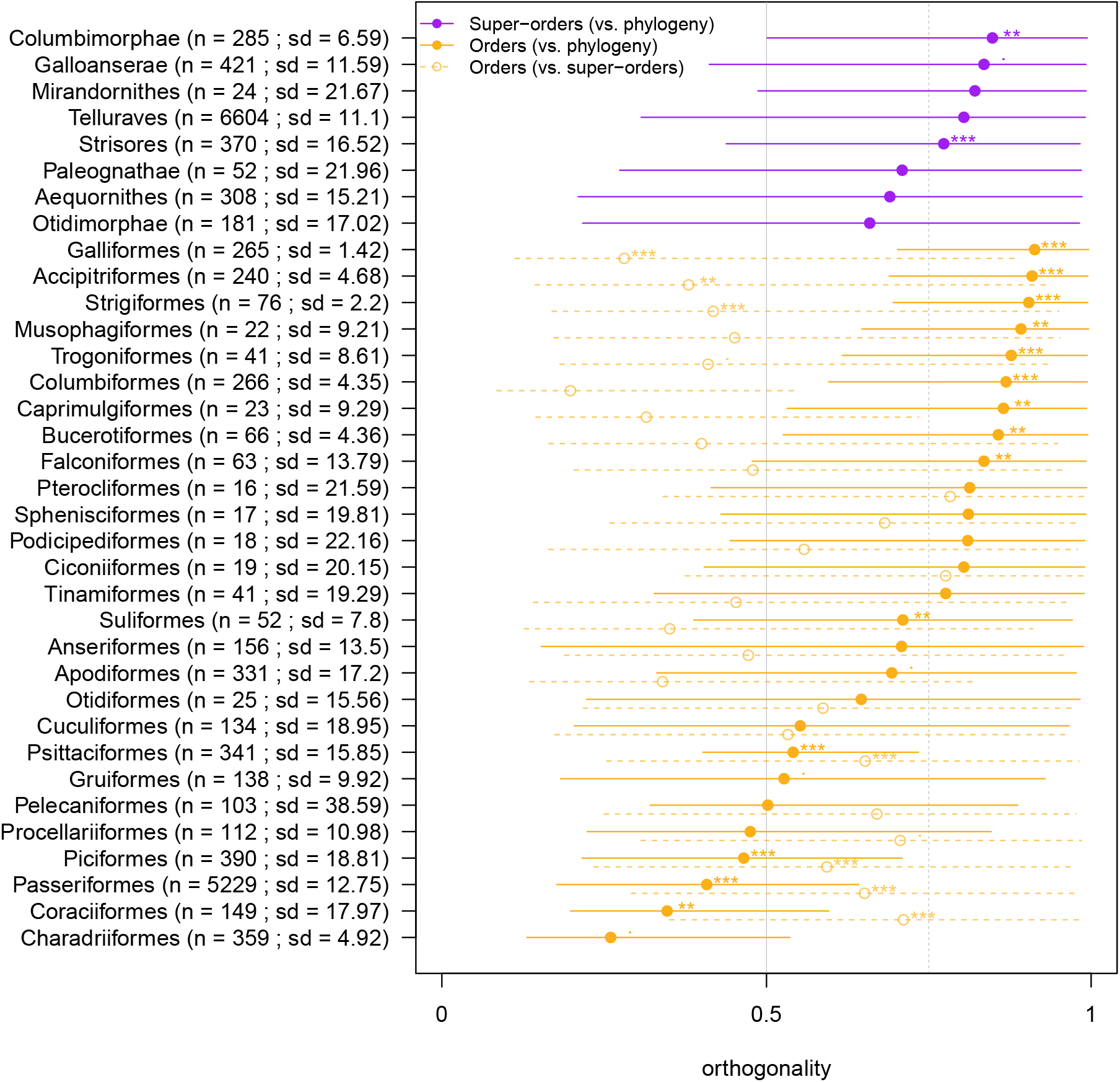
Amount of orthogonality of each clade’s phylogenetic major axis of beak variation compared to their parent or parent’s parent clade. The amount of orthogonality is represented on the horizontal axis and scales from 0 (modulo of 0*°*) to 1 (modulo of 90*°*) with the background gray and dashed gray lines representing, respectively, an orthogonality of 0.5 (modulo of 45*°*) and 0.75 (modulo of 67.5*°*). Dots represent the median orthogonality of each clade and the lines their 95% confidence intervals (CI). No super-order’s phylogenetic major axis of beak variation is parallel to the class-wide phylogenetic major axis of beak variation (purple lines and circles) and 22/27 orders are on average (median) at least half orthogonal (*>* 45*°*) to the class-wide phylogenetic major axis of beak variation. These results are to be contrasted with the orientation of each order relative to their super-order where on average (median) only 14/27 are at least half orthogonal to the orientation of their super-orders’s phylogenetic major axis of beak variation. We also indicate the number of species (n) and the standard deviation (sd) of the orientation of their phylogenetic major axis of beak variation over the 4000 variance-covariance posteriors (sd; expressed in degrees). Note that noclades are clearly parallel to their parent’s or parent’s parent’s ellipses (i.e. no 95% CI includes 0). For each clade we also measured the posterior probability of each clade’s orientation being different from their parent’s clade or the class-wide phylogenetic major axis of beak variation relative to their sample size and sd. The stars represent the posterior probability of the clade’s orientation being different from the comparison clade (*** = pp *>* 0.99; ** = pp *>* 0.95; * = pp *>* 0.9;. = pp *>* 0.8). Only 1/6 super-orders, 8/27 orders relative to class-wide phylogeny and 6/27 relative to their super-orders have a pp *>* 0.99. This is the result of the variation in sample size (n) and standard variation (sd) among clades. A companion plot showing further nested structure within the Passeriformes is shown in Fig. S16.

Substantial variation in *elaboration*_*clade*_ and *innovation*_*clade*_ arises among clades including clades with relatively low *elaboration*_*clade*_ and *innovation*_*clade*_(e.g. Cuculiformes; Fig. 4), low *elaboration*_*clade*_ and high *innovation*_*clade*_ (e.g. Galliformes; Fig. 4), high *elaboration*_*clade*_ and low *innovation*_*clade*_ (e.g. Coraciiformes; Fig. 4), and both relatively high *elaboration*_*clade*_ and *innovation*_*clade*_ (e.g. Podicipediformes; Fig. 4). These patterns hold across scales including within super-orders and at finer scales within the order Passeriformes (Fig. S15). Evolutionary theory predicts that lines of least resistance should break down with time since divergence [14, 15, 21]. We find that the extent of both *innovation*_*clade*_ and *elaboration*_*clade*_ increase with time since divergence from the root of the tree. Similarly, when measured at the macroevolutionary scale, *innovation*_*species*_ and *elaboration*_*species*_ increase with species age (see supplementary Fig. S9 and S10). These results are consistent with both divergence in phenotype that is proportional to time (e.g. as expected under a Brownian motion model) and with the idea that changes in phenotypic correlations are expected to change over longer periods of time, potentially in response to moving adaptive landscapes [24].

Our observations of heterogeneity in the orientation of the phylogenetic major axis of beak variation among clades is further supported by consistent evidence for high amounts of orthogonality of clades relative to the class-wide phylogenetic major axis of beak variation (Fig. 4; Fig. 5; note that these comparisons are based on the evolutionary rate matrix). The median angle of the phylogenetic major axis of beak variation for subclades approaches orthogonality, differing from the class-wide phylogenetic major axis of beak variation by 68.14*°* (95% CI: 22.83*°*-89.09*°*; Fig. 5). Comparisons of orientation among subclades (i.e. orders within super-orders) show similar differences (median = 47.14*°*; 95% CI: 13.64*°*-87.62*°*; Fig. 5), suggesting that reorientations in trait space are largely unconstrained at the megaevolutionary scale and are no more likely to occur along any one axis than another. This differs from previous inference from a subset of our data analysed without incorporating phylogeny, that implied generally consistent and low dimensionality within clades [9]. This suggests that there is no megaevolutionary analogue of the genetic line of least resistance and instead that megaevolutionary shifts more likely reflect changes in the adaptive landscape. Across scales there is remarkable flexibility in the routes to innovation, consistent with the idea that morphological divergence may be less constrained in deep-time than is sometimes assumed [1].

Evolutionary *innovation*_*clade*_ can arise in many directions in trait space (see examples in supplementary Fig. S11). Although the class-wide phylogenetic major axis of beak variation aligns closely with the first dimension of the raw beak shapespace (70.96% of the class-wide phylogenetic major axis of beak variation is aligned with the first dimension of the shape space; Fig. 4), no super-order, and only seven of the 27 orders (Procellariiformes, Pelecaniformes, Charadriiformes, Coraciiformes, Psittaciformes and Passeriformes), are aligned mainly PC1 (Fig. 4-bar plots); *innovation*_*clade*_ dominates at this scale. This pattern of orthogonality also holds for sub-orders and families within the Passeriformes where only half of the sub-orders (Meliphagoidea, Corvides and Passerida) and seven of the 23 families (Eurylaimides, Meliphagoidea, Petroicidea, Fringilidae, Aegit-haloidea, Pyconotiae, Nectariniidae) align with the Passeriformes phylogenetic major axis of beak variation (Fig. S15). In addition to the lack of alignment of clade phylogenetic major axis of beak variation with the class-wide phylogenetic major axis of beak variation for birds, we also found that within clades, beak shape variation is either highly constrained (varying almost entirely along a single axis; e.g. in Galliformes) or higher-dimensional than in some orders or superorder than across the whole class Aves (for example, there is no clear dominant phylogenetic major axis of beak variation; e.g. in Podicipediformes) from the class-wide phylogenetic major axis of beak variation. For example, Accipitriformes (hawks and allies) have phylogenetic major axes of beak variation that mainly align with the second dimension of the shapespace suggesting distinct directions of beak evolution for the clade relative to the class-wide phylogenetic level but uniformity in their beak shape within the clade. In contrast, the Podicipediformes (grebes) are highly variable across all dimensions suggesting that all components of the shapespace are necessary to describe their beaks. These observations illustrate multiple routes to *innovation*_*clade*_ and imply that within clades, species beak shapes can evolve in many directions (akin to Endler *et al*.’s [46] random innovation) or follow a single direction (akin to Endler *et al*.’s [46] specific innovation) and that reorientations of trait space can arise in any direction, in any lineage, and at any time throughout avian history.

### 3.4 Species elaborate *and* innovate, clades innovate

The disconnect between observations of reorientation of trait space at the macroevolutionary scale, and an apparent predominance of *elaboration*_*species*_ could be viewed as a paradox. We formalize this paradox by testing whether elaboration is more common at higher taxonomic levels (megaevolutionary scale) than at the species level (macroevolutionary scale). We compared the area under the curve of the scaled density of *elaboration*_*species*_, *innovation*_*species*_, *elaboration*_*clade*_ and *innovation*_*clade*_. We then measured the difference between both areas and found that there was a significant difference in *innovation*_*species*_ and *innovation*_*clade*_ but no clear difference in *elaboration*_*species*_ and *elaboration*_*clade*_ (Fig. 6). This difference confirms our observation that innovation is indeed common and dominant at the megaevolutionary scale, however, the contributions of elaboration and innovation are shown to be indistinguishable at the macroevolutionary scale. So, although species often evolve preferentially along a shared phylogenetic major axes of beak variation within clades, there is frequent deviation from these major axes, and the phylogenetic major axes of beak variation change among clades. This could be explained in part because the evolutionary rate matrix (**R**) is more influenced by shifts in the adaptive landscape through time and thus reflecting natural selection in deeper nodes more than in younger nodes (Fig. S9 and S10).

**Figure 6:**
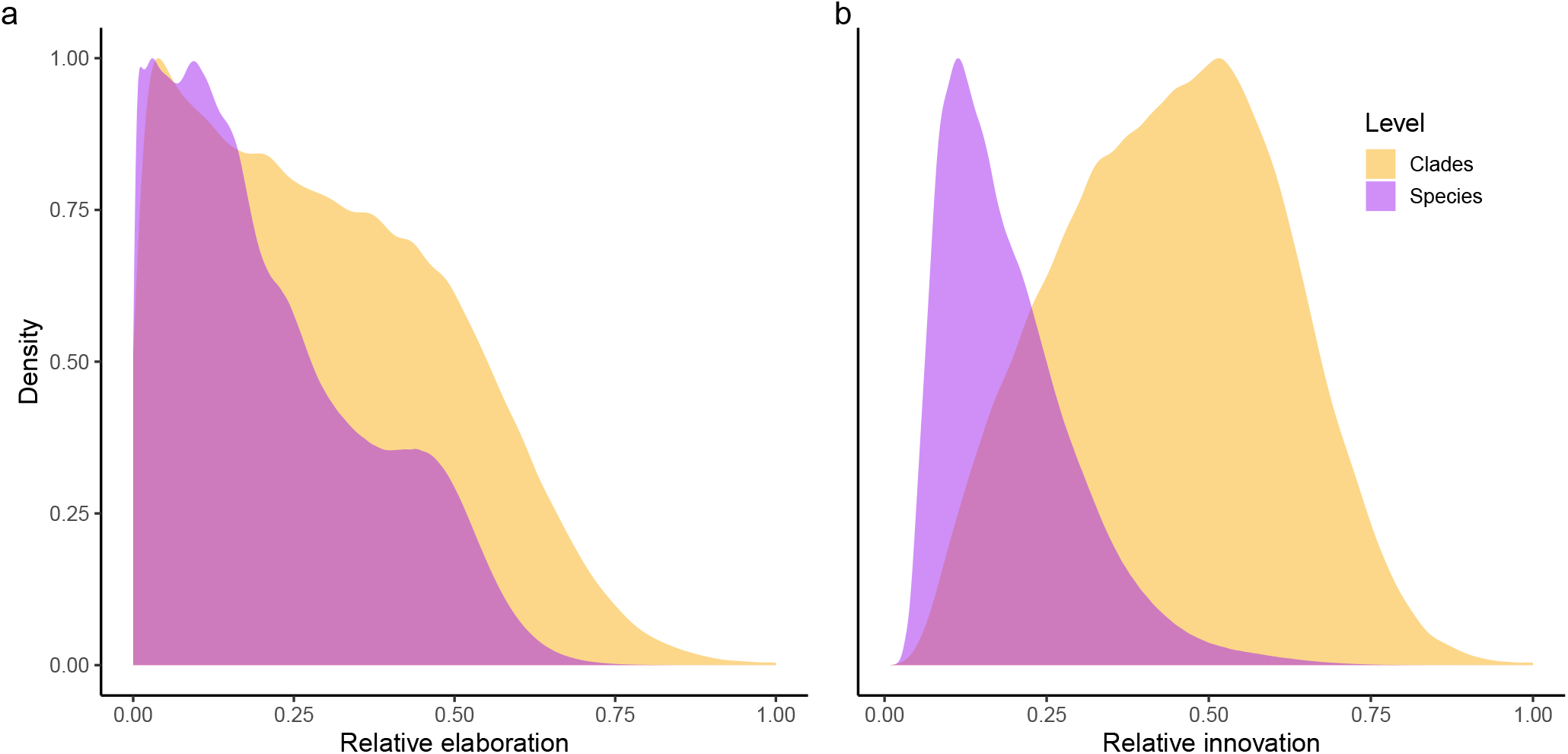
Comparisons of relative *elaboration*_*species*_ and *innovation*_*species*_ (Fig. 2) and *elaboration*_*clade*_ and *innovation*_*clade*_ (Fig. 4). a) the *elaboration*_*species*_ (purple) for each species in each 4000 posterior samples compared to the *elaboration*_*clade*_ for each of the 35 clades (yellow). The *elaboration*_*clade*_ scores are scaled by the maximum score within the clade (corresponding to the scaled ellipses in Fig. 4) and the scores for species are scaled by the maximum *elaboration*_*species*_ score. This allows us to compare *elaboration*_*species*_ in and *elaboration*_*clade*_ despite their different sample sizes. We use the Bhattacharryya Coefficient to quantify the overlap of the area under the curve for species and clades. For elaboration the overlap is Bhattacharryya Coefficient = 0.217, indicating no distinguishable dissimilarity in the amount of *elaboration*_*species*_ and *elaboration*_*clade*_ across the posterior samples. b) as a) but using *innovation*_*species*_ and *innovation*_*clade*_ scores. For innovation the overlap has Bhattacharryya Coefficient = 0.039, which indicates a clear difference in the amount of *innovation*_*species*_ and *innovation*_*clade*_ across the posterior samples.

We further tested the contributions of elaboration and innovation at different scales by examining the consequences of elaboration and innovation for the observed divergence of bird beak shapes. Specifically, we calculated the distance to centroid of avian beak morphospace for each species (Fig. 2). We then used phylogenetic generalised least squares (PGLS; [52]) to model distance to centroid as a function of *elaboration*_*species*_, *innovation*_*species*_ and *elaboration*_*clade*_ and *innovation*_*clade*_. We ran the PGLS analysis using both single and multiple predictors. Our analyses indicate that the majority of variation (r^2^ = 0.888; Table 1) in beak shape centroid distance can be explained by a combination of *elaboration*_*species*_ and *innovation*_*species*_. When modeled as single predictor models, variation explained is lower for *elaboration*_*species*_ than *innovation*_*species*_ and *innovation*_*species*_ has a steeper slope indicating that although elaboration is overall the dominant model of divergence, innovation leads to greater exploration of morphospace (Table 1). In contrast, the combination of *elaboration*_*clade*_ and *innovation*_*clade*_ alone explains only *>*0.01% of the total variation in beak shape divergence (Table 1). However, *innovation*_*clade*_ is nonetheless an important contributor to total beak shape space. We reach this conclusion because models with interactions between species and clade metrics provide by far the best model fit overall and removal of terms indicate that the most important interactions are between *innovation*_*clade*_ and *innovation*_*species*_ followed by *innovation*_*clade*_ and *elaboration*_*species*_. The interaction terms show that the relationship between species metrics (*elaboration*_*species*_, *innovation*_*clade*_) and distance to centroid becomes steeper as *innovation*_*clade*_ increases (Table 1). Overall, the models suggest that although most species evolve via elaboration at the macroevolutionary scale, expansions of beak morphospace are also driven by megaevolutionary scale reorientations of trait space.

**Table 1:** Elaboration and innovation as predictors of beak shape distance from centroid. The table shows parameter estimates of predictors from 12 different phylogenetic generalised least squares (PGLS) models with ΔAIC showing relative model fit following removal of terms. Models 1-5 are multiple regressions including all variables and one or more interaction terms (e.g. model 1 contains all the terms and their interactions); models 6-8 are are multiple regressions including two or more variables and no interaction terms (e.g. model 6 contains all the terms but no interactions); models 9-12 are single predictor models (e.g. model 9 contains only one term). Values in the table are based on PGLS in which Pagel’s *λ* was fixed at 1 (i.e. assuming a Brownian motion model of evolution). Fixing *λ* is necessary to allow model comparison with AIC. We also fitted the same models fixing *λ* to 0.727 (the lowest value of *λ* from any individual model) and relative model fit and interpretation of parameters was unaffected. *E* = elaboration; *I* = innovation; AIC = Akaike Information Criterion; *** = p-value *<* 0.001.

| Model | $E_{species}$ | $I_{species}$ | $E_{clade}$ | $I_{clade}$ | $E_{species} \times I_{clade}$ | $E_{species} \times E_{clade}$ | $I_{species} \times I_{clade}$ | $I_{species} \times E_{clade}$ | $\Delta AIC$ | Adj. $r^2$ |
| --- | --- | --- | --- | --- | --- | --- | --- | --- | --- | --- |
| 1 | 0.04*** | -0.139*** | 0.002 | -0.002 | 0.064*** | 0.045*** | 0.341*** | 0.093*** | 0.00 | 0.92 |
| 2 | 0.257*** | 0.169*** | 0.01 | 0.001 | 0.045*** |  | 0.312*** | 0.034*** | -691.48 | 0.92 |
| 3 | 0.108*** | 0.236*** | 0.012 | -0.002 | 0.055*** | 0.034*** | 0.343*** |  | -284.74 | 0.92 |
| 4 | 0.254*** | 0.104*** | 0.002 | 0.004 |  | 0.027*** | 0.311*** | 0.052*** | -877.85 | 0.92 |
| 5 | 0.1*** | 0.571*** | -0.018* | 0.02*** | 0.054*** | 0.036*** |  | 0.096*** | -1498.36 | 0.91 |
| 6 | 0.364*** | 0.973*** | -0.007 | 0.023*** |  |  |  |  | -2230.10 | 0.90 |
| 7 | 0.364*** | 0.972*** |  |  |  |  |  |  | -2280.08 | 0.90 |
| 8 |  |  | -0.025 | 0.011 |  |  |  |  | -22270.55 | -0.00 |
| 9 | 0.286*** |  |  |  |  |  |  |  | -18986.63 | 0.32 |
| 10 |  | 0.798*** |  |  |  |  |  |  | -17714.99 | 0.41 |
| 11 |  |  | -0.008 |  |  |  |  |  | -22269.34 | -0.00 |
| 12 |  |  |  | 0.002 |  |  |  |  | -22269.51 | -0.00 |

Taken together, these results suggest that rather than a paradox, the reorientation of trait space can instead arise as a result of species-level innovations within clades arising along common directions in phenotypic space. This idea is similar in concept to multiple adaptive peak models where species evolve in a heterogeneous adaptive landscape and share similar responses to selection pressures within adaptive zones [53]. This implies that gradual directional evolution [6], rather than exceptional megaevolutionary jumps [1, 9], may be sufficient to explain diversity in avian beak morphology. In other words, populations can always change morphologically within the shapespace with minimal amounts of innovation over long periods of time.

## 3.5 Conclusions

Our results show that although at the macroevolutionary scale most bird beaks are elaborating along a class-wide or sub-clade phylogenetic major axis of beak variation, at the megaevolutionary scale, innovation away from the class-wide phylogenetic major axis of beak variation is much more common. This nested structure of elaboration at a lower (species) taxonomic level and innovation at a higher (clade) taxonomic levels could thus explain the diversity of bird beaks we observe today. However, individual species-level variation in the past could also have led to a shift in a clade’s line of least evolutionary resistance through different evolutionary routes on a dynamic adaptive landscape. Taken together, our results suggest that the signature of evolutionary reorientations in deep-time (*innovation*_*clade*_), coupled with *elaboration*_*species*_, is a robust explanation of the massive diversity of bird beaks we observe today and is consistent with recent suggestions from univariate analysis that observation of apparently abrupt phenotypic shifts can be explained by gradual Darwinian processes [6, 24, 11]. Indeed, the class-wide phylogenetic major axis of phenotypic variation is an emergent property of reorientation of trait space among clades that requires no special evolutionary process: megaevolutionary patterns appear to be an inevitable outcome of evolution on a shifting adaptive landscape.

## 4 Materials and Methods

### Beak shape data

We used the dataset from [9, 47, 54] which consists of four landmarks and three curves of 25 semi-landmarks each, taken from 3D scans of the beaks of 8748 bird species. The landmark data were aligned with Procrustes superimposition using the R package `Morpho` [55, 56] and we removed the size component and controlled for symmetry (see [9, 47, 54] for detailed descriptions of data collection and processing). We ordinated the superimposed landmarks using a principal components analysis (PCA) and selected the eight first dimensions of the resulting PCA. These eight dimensions contained at least 95% of variance in each super-order, and 98.7% of the variance in the whole dataset. We used a threshold of at least 95% of variance explained for each super-order because although the overall distribution of the variance on each PC axis in the whole trait-space is decreasing (i.e. *s* PC1 *> s* PC2 *> s* PC3), this is not always true for each super-order. For example, in the Mirandornithes, PC2, then PC4, and then PC1 contained the most variance (Fig. 4 and S2). This protocol for selecting the number of PC axes ensures that the resulting trait-space contains enough variance for each super-order (see Fig. S2). This procedure resulted in an 8748 × 8 matrix that contains at least 95% of the variance in all observed living bird beaks (hereafter, the shapespace).

### Phylogenetic data

We used a random subsample of 1000 trees from the avian tree distribution of [57] for the analyses and one randomly selected tree for the figures (Fig. 2). We pruned these trees to contain only the 8748 species present in our beak shapespace.

### Phylogenetic GLMM analysis

We ran a Bayesian phylogenetic generalised linear mixed model (pGLMM) using the `MCMCglmm` and `ape` R packages [48, 58] with the shapespace data as a multidimensional response variable, the error in the dataset as residuals, and each clade’s phylogeny as a nested random term, i.e., a model of:

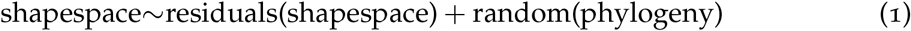

This model allows us to estimate an evolutionary rate matrix, here defined as a matrix containing the rate of among-species divergence in species traits that explicitly incorporates phylogeny (**R** matrix of [28] and **B** matrix of [29])) for each clade and the phylogeny as a whole, and one for the residuals in the shapespace itself [50]. These evolutionary rate matrices were then used in a base projection-rejection analysis to measure the elaboration and innovation scores (see below). We ran two separate nested models. The first model used one random term for the class-wide phylogeny and 35 nested random terms, one for each super-order and order containing more than 15 species in our dataset (Fig. 4). This resulted in a multidimensional GLMM with a 8748 × 8 response variables, one class-wide residual term and 36 random terms. The second model was fitted to a subset of the dataset containing only the 5229 Passeriformes species with 30 nested random terms, one for each sub-order and families containing more than 15 species in our dataset (Fig. S15. This resulted in a multidimensional GLMM with a 5229 × 8 response variables, one order-wide residual term and 30 random terms. To account for phylogenetic uncertainty we ran the model using different tree topologies from the tree distribution in [57]. Because of the very large size of our model and dataset, however, running the full model on multiple trees was computationally unfeasible (it would require 2 CPU years and 4TB of RAM for each of the models, i.e. the class-wide phylogeny and the Passeriformes models). Instead, we developed and implemented a “mini-chains” MCMCglmm method using the R package `mcmcmglmmm` [59].

### mcmcmcglmmm

This method runs relatively short Monte-Carlo Markov chains (MCMC) across a large number of different phylogenetic trees and concatenates the resulting short posteriors into one larger posterior distribution that contains the phylogenetic uncertainty information. We performed the method using the following protocol (see Fig. S1). First we ran three independent MCMCglmm chains, hereafter the parameterisation chains, for 50,000 generations with the trait data and the model described above on the consensus tree from the tree distribution, along with flat priors with a belief parameter of 2%, i.e. with a very low weight on the priors. Next we extracted the following parameters from the parameterisation chains: a) the conservative burnin average time across the three chains, defined as the highest number of iterations required to first reach the median posterior likelihood value multiplied by 1.1; and b) the mini-chains priors, defined as the median posterior model values from the parameterisation chains with a belief parameter of 5%. We then ran 400 mini chains in parallel with the estimated burnin time and priors to run 10 samples past the burnin. This resulted in 10 exploitable posterior samples for each tree. We used 400 trees because that was the number of models required to reach an effective sample size (ESS) of at least 200 for all the estimated parameters (see Fig. S3 and SS4). Using this approach, we reduced the computational time to around 45 CPU hours and 9GB of RAM per mini-chain - a 400 fold improvement. The total analysis took around three CPU years using the shARC cluster from the University of Sheffield. Code to reproduce the procedure is available in the `mcmcmcglmmm` vignette [59].

### Elaboration/innovation scores using projection/rejection analyses

We used the distributions of the 4000 posterior evolutionary rate matrices to run the projection/rejection analyses to obtain elaboration and innovation scores for bird beaks. We used linear algebra to interpret elaboration and innovation *sensu* [46] by using the major axis of the evolutionary rate matrices (referred to throughout the text as the phylogenetic major axis of beak variation) as the “line of least resistance”, or, specifically here, the line of elaboration for each random term (corresponding to the elaboration axis for each clade). We projected either species or another phylogenetic major axis of beak variation onto that clade. We can then interpret where a species or a clade falls on the phylogenetic major axis of beak variation of reference as its *elaboration*_*species*_ score, and how far a species is from that phylogenetic major axis of beak variation as its *innovation*_*species*_ score. We calculated the elaboration and innovation scores using two main approaches: 1) by clades (i.e. to measure the *elaboration*_*clade*_/*innovation*_*clade*_ at the megaevolutionary scale)where we calculated the projections of the phylogenetic major axis of beak variation of each clade onto the class-wide phylogenetic major axis of beak variation (Fig. 4); and 2) by species (i.e. to measure the *elaboration*_*species*_/*innovation*_*species*_ at the macroevolutionary scale) where we calculated the projections of each species onto a) the class-wide phylogenetic major axis of beak variation and b) the phylogenetic major axis of beak variation of their parent clade (Fig. 2). To make the results easier to interpret, we centered and scaled elaboration and innovation scores onto the center of the phylogenetic major axis of beak variation of interest and made the elaboration values absolute. In this way, we can interpret an elaboration or innovation score within the 0-1 range to be non exceptional, i.e within the 95% confidence interval of the evolutionary rate matrix. The full mathematical procedure is described in detail in the supplementary materials (section 4; note that the procedure described is applied to each individual evolutionary rate matrix for each term in the model, e.g. for each 4000 posterior variance matrices individually for each *elaboration*_*species*_, *innovation*_*species*_, *elaboration*_*clade*_ or *innovation*_*clade*_ score). Code to reproduce the analyses is available in this `dispRity` vignette.

### Clade orthogonality

We measured the amount of orthogonality of each random term (i.e. clade) in our models compared to their parent clade and their parent’s parent clade. For example for Columbiformes, we measured the amount of orthogonality in the evolutionary ratematrices of (i) the Columbimorphae random terms against the phylogeny random terms, (ii) the Columbiformes random terms against the Columbimorphae random terms (parent) and (iii) the Columbiformes random terms the whole phylogeny random terms (parent’s parent). To measure the amount of orthogonality we calculated the angles between the phylogenetic major axis of beak variation of the focal clade and their parent clade for each posterior evolutionary rate matrix and within each clade, by randomly comparing phylogenetic major axes of beak variation within a clade. We converted each angle measurement into a degree of orthogonality metric where 0 corresponds to flat angles (0*°* or 180*°*) and 1 to right angles (90*°* or 270*°*). We then measured the posterior probability of the angles within a clade being different to the angles among clades. These results are available in Fig. 5. Note that we did not use the random skewers method here [60] because we were interested in measuring the amount of orthogonality between pairs of evolutionary rate matrices rather than simply measuring whether they were different (but see supplementary materials Fig. S13 for random skewers correlation results).

### Phylogenetic generalised least squares analysis

We ran phylogenetic generalised least squares models (PGLS) to test the effects of elaboration and innovation on beak shape divergence. Beak shape divergence was calculated as the Euclidean distance of each species beak shape from the centroid of beak shape of all birds. We then fitted a series of nested models with distance as the response and *elaboration*_*species*_, *innovation*_*species*_, *elaboration*_*clade*_, and *innovation*_*clade*_ as predictors. The most complex model included all two-way interaction terms:

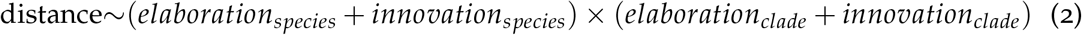

We then sequentially removed terms to examine effects on model fit. We initially estimated Pagel’s *λ* [61] for all models. However, model comparison based on AIC scores can be misleading when estimating *λ* (i.e. model differences can arise because of different *λ* fits, rather than due to effects of the parameters of interest). We therefore repeated all models a further two times, once fixing *λ* to 1 (equivalent to assuming Brownian motion model of evolution) and once fixing *λ* to the lowest value estimated from any of the models. The results were qualitatively similar regardless of *λ* and we report only models based on pure Brownian motion in the main text. In addition, we estimated Pagel’s *λ* for *elaboration*_*species*_ and *innovation*_*species*_ as univariate traits. All models were fitted using the phylolm library using a single phylogeny [52].

## Reproducibility and repeatability

The raw and processed data used in this analysis is available from figshare [62]. The code used to reproduce the figures and tables is available from github [63].

## Acknowledgements

We thank M. Adams, H. van Grouw and R. Prys-Jones from the Bird Group at the NHM, Tring and H. McGhie at the Manchester Museum for providing access to and expertise in the collections. We are indebted to the volunteer citizen scientists at http://www.markmybird.org for helping to build the database of bird bill shape and contribute to our understanding of avian evolution. We thank Masahito Tusboi and three other anonymous reviewers for their suggestions that greatly helped improving the manuscript.

## Author contributions

G.H.T., A.P.B., N.C., and T. G. designed the study. J.A.B., C.R.C., E.C.H., Z.K.V., and G.H.T. curated the data. T.G. analysed the data. All authors contributed to the writing of the manuscript.

